# Thermoplastic polyurethane as a base material for soft, high-density neural interfaces

**DOI:** 10.64898/2026.10.04.755411

**Authors:** Anna Pak, Andrada Iulia Velea, Tim Hosman, Joshua Wilson, Raphael Panskus, Manuel Seckel, Pablo de Anta Dardagan, Ralf Schmidt, Domonkos Horváth, Kinga Tóth, Domokos Meszéna, Astrid Gollhardt, Nasim Bakhshaee Babaroud, Barbara Pahl, Eleni Zingkou, Asimina Kolianou, Georgia Sotiropoulou, Sandeep Negi, Florian Solzbacher, Christine Kallmayer, István Ulbert, Vasiliki Giagka

## Abstract

Polymers used as substrate and encapsulation for implants are typically deposited layer-by-layer, leaving polymer–polymer interfaces that adhere poorly and become a critical failure point. Thermoplastic polyurethane (TPU) elastomers combine four properties not found together in any other candidate: decades of clinical use in long-term implants and availability in medical grade, a stiffness well below that of polyimide and parylene C, compatibility with microfabrication, and thermoplastic self-fusion, which removes the polymer–polymer interface entirely. This work introduces TPU as a base material for soft, high-density neural interfaces and demonstrates one route to realizing them. Gold is patterned on a sacrificial copper sheet by maskless lithography and transferred into TPU by vacuum lamination, giving conformally coated tracks down to 15 µm at 30 µm pitch within a single, merged polymer body. An additional electroplating step produces a nanostructured “pinecone” gold surface that increases the electrode electrochemically active area, approximately doubling the charge injection capacity. Daisy-chain silicon chips were also embedded within the soft polymer body. The films transmit up to 88% at 470 nm, a wavelength commonly used for optogenetics. TPU showed no cytotoxicity towards differentiated SH-SY5Y cells and only a mild tissue response after 9.5 months of subcutaneous rat implantation.

## 1 Introduction

Electrical stimulation of active cells by means of neural implantable devices is a powerful tool for treating various neurological disorders [1–3]. When combined with electrical recording, it can provide a deeper understanding of the underlying neurophysiological processes [4, 5]. Neural interfaces can be made on rigid or soft/flexible substrates. Rigid substrates, typically silicon (Si), are broadly compatible with common microfabrication techniques, leading to small feature sizes and high resolution [6–10]. However, they leave a larger impact on the tissue and cause a stronger foreign body reaction (FBR) [11–13]. Polymer-based neural interfaces, which more closely match the mechanical properties of the tissue, are thus often preferred. Polyimide, parylene and silicone-based neural interfaces have demonstrated great tissue compatibility in a variety of applications [14–16]. Even among polymers, there is evidence suggesting that softer materials, e.g., silicone elastomers, minimize the impact the implant has on soft tissue compared to stiffer, even though flexible, counterparts, e.g. polyimides [16].

However, silicones are not easily compatible with microfabrication techniques, hence silicone-based neural interfaces suffer from limited feature density and resolution [17, 18]. Further, most biocompatible polymers commonly used as neural implant substrates and coatings, are deposited in a layer-by-layer fashion using chemical vapour deposition (CVD) or spin coating [15, 19–23]. Because of this layer-by-layer deposition, these polymers face the problem of poor adhesion and delamination at the polymer-polymer interface [24–28]. A number of strategies, such as plasma and corona discharge pre-treatment [29], annealing [26], adhesion promoters [27], interlayers [14], and partial curing [28] are used to improve the adhesion between these polymer layers. Nevertheless, the interface always remains present and is considered a critical point of failure, compromising the long-term stability, and thereby functionality, of the implant in the aqueous body environment [24]. On the other hand, thermoplastic polymers, such as liquid crystal polymer (LCP), have demonstrated long-term stability in wet environments and an absence of delamination, due to the complete merging of the polymer with itself after thermocompression bonding [30–32]. However, LCP is relatively rigid, opaque, and requires high temperatures for processing (around 300°C), limiting its range of applications for neural interfaces.

Thermoplastic polyurethane (TPU) is a thermoplastic elastomer, a segmented block copolymer whose alternating hard and soft domains give rubber-like elasticity while the material remains melt-processable [33]. It is, to our knowledge, the only candidate that combines four properties relevant to chronic neural interfaces. First, polyurethanes have several decades of clinical use in long-term implants, as insulation for pacemaker leads, in vascular and intravenous catheters [34, 35], and vascular grafts [36], and are available in medical grade (USP Class VI) [37]. Second, they are considerably softer than polyimide and parylene-C, closer in Young’s modulus to silicone elastomers and soft brain tissue (Figure 1a). Third, unlike silicone elastomers, TPUs are compatible with microfabrication and support high-resolution patterning. Fourth, being thermoplastic, TPU merges with itself under heat and pressure, removing the polymer–polymer interface rather than attempting to strengthen it. Its transparency additionally permits optical access for imaging and optogenetic stimulation. Polyurethane has appeared in neural devices in other roles: blended with PEDOT:PSS as a stretchable conductive composite [38], and in bioresorbable, transient constructs [39], while our own earlier work addressed individual process steps [40] and the acoustic properties of the material stack [41, 42], without functional interfaces. To the best of our knowledge, TPU has not previously been used as the substrate and encapsulation material of a micro-fabricated, thin-film metallised neural interface. Finer features have been achieved on photopatternable fluorinated elastomers [43], which are crosslinked rather than thermoplastic and therefore retain an interface between successive layers.

**Figure 1:**
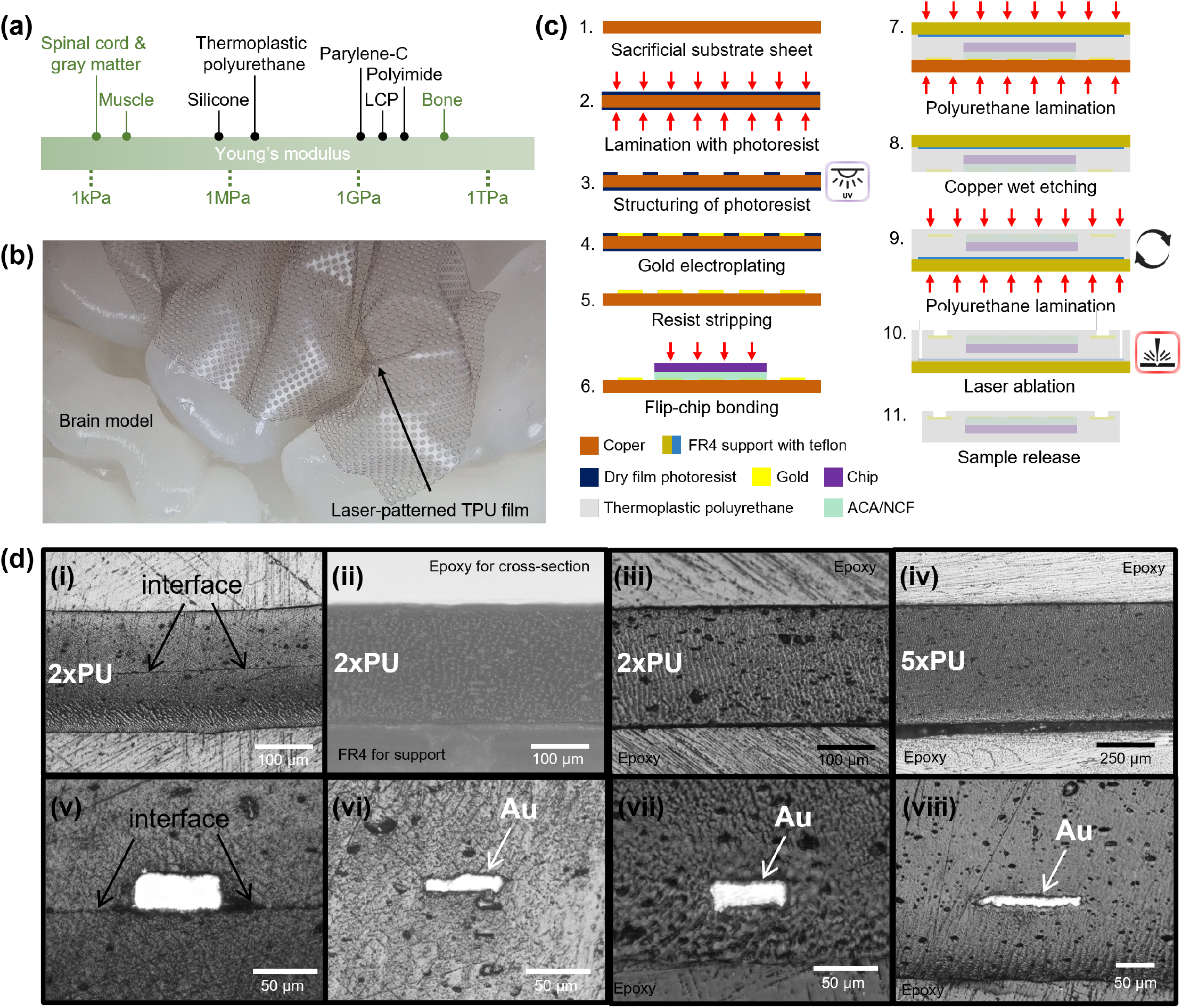
(a) Mechanical mismatch in Young’s modulus between body tissues and commonly used polymers for neural implants. Inspired by [23]. Young’s modulus ranges, collected from [44–54]. (b) A laser-micropatterned TPU film draped over a gyrified brain model, following the curvature of the gyral crowns and shallow inter-gyral valleys. (c) Step-by-step schematic cross-section representation of the fabrication process. Flip-chip bonding step is optional to achieve active devices. (d) Cross-section image of two non-merged TPU sheets with existing interface laminated under non-optimised parameters (i); cross-section of two completely merged TPU sheets directly after lamination (ii) and after 100 days soak in PBS at 37°C (iii); cross-section of five completely merged TPU sheets after six months of implantation (iv). 2*−* TPU and 5 *−*TPU indicating two and five merged TPU layers. Cross-section of two non-merged TPU sheets around the gold track with existing interface laminated under non-optimised parameters (v); cross-section of two completely merged TPU sheets with embedded gold track directly after lamination (vi) and after 100 days soak in PBS at 37°C (vii); cross-section of five completely merged TPU sheets around the gold track after six months of implantation (viii). 100 µm thick sheets of Platilon U 4201 AU TPU were used.

Here, the thermoplastic properties of polyurethane are used to create a homogeneous, thin, soft, and flexible insulating environment within which dense metallization and small, thin chips can be embedded, enabling high-density and high-resolution, soft, and, eventually, active, implantable neural interfaces. We verify the suitability of TPU for use in neural implants using both *in vitro* cytotoxicity tests on differentiated SH-SY5Y neuroblastoma cells and *in vivo* tests in rats. We employ a polymer-last approach, which, coupled with laser direct imaging (LDI)-based photolithography, enables the creation of dense, high-resolution gold (Au) patterns without photomasks, yielding an easily adjustable, inexpensive process. We use flip-chip bonding to demonstrate the integration of active components and illustrate the technology’s versatility by creating prototypes of neural implants with dummy chips or other silicon-based components, with form factors suitable for interfacing with the peripheral and the central nervous system. This result, combined with the availability of medical grade TPU with excellent mechanical properties, makes the proposed approach a prime candidate for the development of translational neural implants that have the ability to conform to soft neural tissue.

## 2 Results

### 2.1 Softness and conformability of TPU

TPU is a thermoplastic elastomer, and its Young’s modulus places it between the silicone elastomers and the polymers conventionally used for high-density neural interfaces (Figure 1a). For a thin film, conformability is governed by the bending stiffness (*D* = *Et*^3^*/*[12(1 − *ν*^2^)], where *t* is the film thickness and *ν* the Poisson’s ratio) rather than by the modulus (*E*) alone. At equal thickness, a TPU film is approximately two orders of magnitude less stiff in bending than a polyimide one, corresponding to roughly a tenfold reduction in the minimum radius it can follow. Following a doubly curved surface such as the cortex additionally requires in-plane deformation. The TPU films were therefore micropatterned by picosecond laser machining with a regular array of through-holes (Figure 1b), allowing local accommodation of in-plane strain. The patterned films follow the curvature of the gyral crowns and of the shallow valleys between them, at the cost of reduced continuous surface area.

### 2.2 Fabrication process

The fabrication process for embedding the metal tracks and the silicon chip in soft polyurethane is schematically illustrated in Figure 1c.

In short, a copper (Cu) sheet is used as a sacrificial substrate and coated with dry-film photoresist. LDI photolithography is used to expose the Cu areas that will later form tracks, electrodes and pads for subsequent Au electroplating, after which the resist is stripped chemically. At this point, a thin silicon chip can be connected to the pads by flip-chip bonding using an adhesive/underfill (anisotropic conductive adhesive (ACA) or TPU as non-conductive film (NCF)); passive arrays are produced by omitting this step. The first TPU layer, either a polyether TPU (Platilon U 4201 AU, Covestro) or a medical-grade polycarbonate-urethane (Bionate 80A, dsm-firmenich), is then laminated on top and the sacrificial Cu sheet is etched away. The stack is then flipped, and the second TPU layer is laminated, embedding the metallization within a single, merged polymer body. Electrode and pad openings are finally made by laser ablation. Full details are given in the Experimental Section. The sample configurations used throughout this work are summarized in Table S1.

### 2.3 Fusion of polyurethane layers and conformal coverage of gold tracks

The central requirement of the proposed approach is that the two laminated TPU layers merge into a single, continuous polymer body, leaving no polymer–polymer interface at which delamination could initiate. Whether this occurs depends on the lamination parameters. When two 100 µm thick TPU sheets were laminated under non-optimized conditions (vacuum lamination at 145 °C and 100 N cm^*−*2^), the original interface remained clearly visible in the cross-section as a continuous line running through the stack (Figure 1d-i, with the lamination profile shown in Figure S2b). After optimization of the lamination profile (vacuum lamination at 165 °C and 100 N cm^*−*2^; profile in Figure S2c), no interface was visible between the two layers, which appeared as a single homogeneous body (Figure 1d-ii). The same behavior was observed around the metallization. Under non-optimized parameters, the interface between the two layers remained visible where they met around an embedded Au track (Figure 1d-v). After optimization, the tracks were conformally covered, with the polymer following the track profile (Figure 1d-vi). Both the merged layers and the conformal coverage persisted under prolonged exposure to physiological conditions. After 100 days of soaking in PBS at 37 °C, no interface had reappeared in the merged stack (Figure 1d-iii), and the embedded gold track remained conformally covered (Figure 1d-vii). The same was found after six months of subcutaneous implantation, for the polymer stack and around the gold track (Figure 1d-iv,viii, cross-section of, in this case, 5 TPU layers), indicating that the merged layers remain stable *in vivo* over this period.

### 2.4 Electrode characterization

Two types of electrodes were fabricated: “cauliflower” electrodes (Figure 2a), obtained directly after laser ablation, and nanostructured “pinecone” electrodes (Figure 2b), obtained through an additional, modified electroplating step after laser ablation (Section 5). Scanning electron microscopy (SEM) images of the resulting electroplated surfaces of the “cauliflower” and “pinecone” electrodes are shown in Figure 2a and Figure 2b, respectively, and show that the two surfaces differ markedly in morphology. The cauliflower-like electrodes appear to exhibit a granular, agglomerated microstructure with clustered, rounded features, whereas the pinecone-like electrodes display a hierarchical micro-nanostructure with a branched, lamellar morphology, consisting of sharp, flake-like layers.

**Figure 2:**
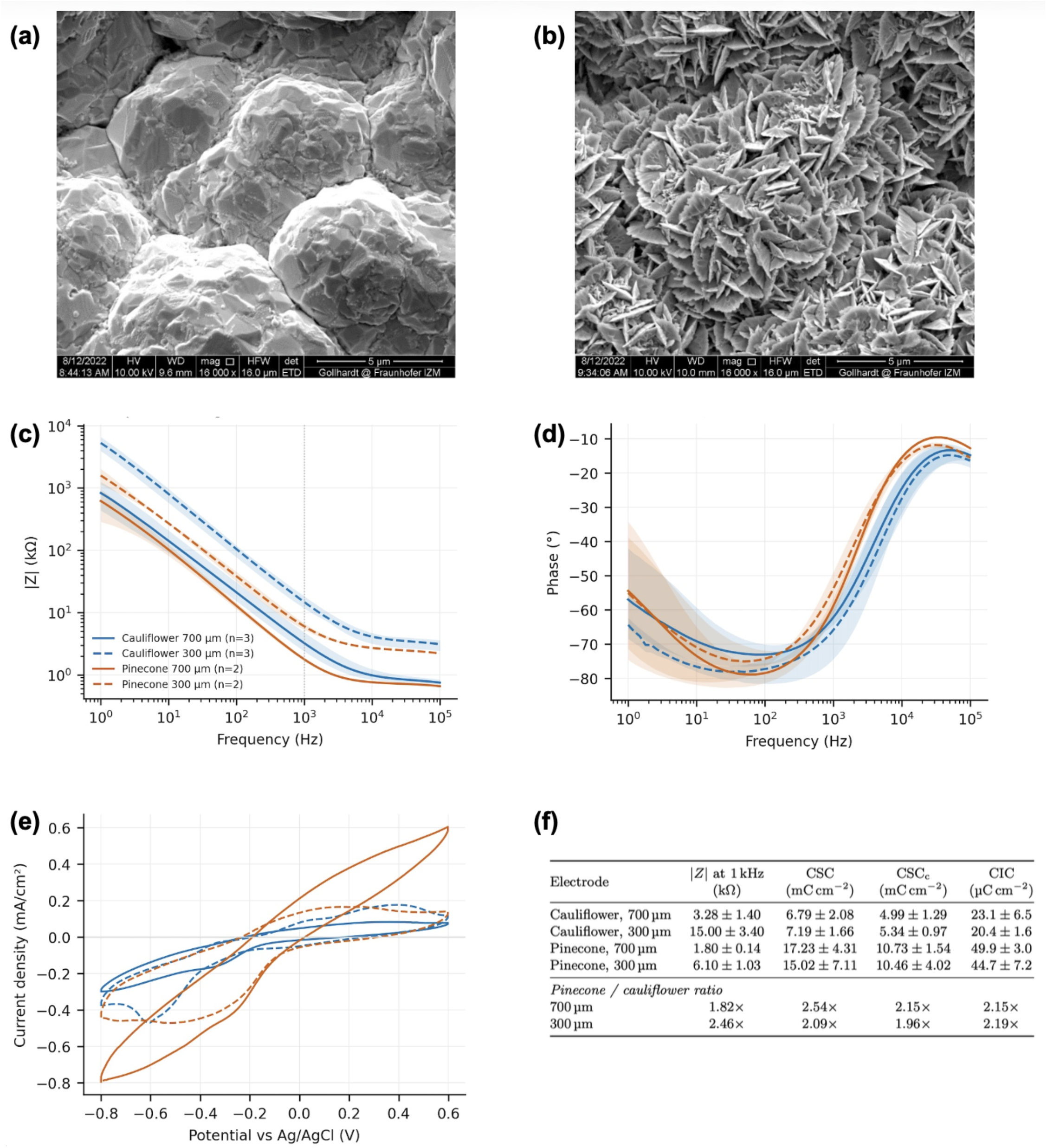
(a-b) SEM images of (a) “cauliflower” and (b) nanostructured “pinecone” electroplated gold. (c) Impedance magnitude, (d) phase angle plots and (e) cyclic voltammograms (50 mV/s scan rate), for 700 and 300 µm “cauliflower” and “pinecone” electrodes; lines are the mean and shaded bands the range across electrodes. (f) Electrochemical characterization summary: Impedance magnitude at 1 kHz, total and cathodic charge storage capacity (CSC, CSC_c_) and maximum cathodic charge injection capacity (CIC) for the four electrode types. Electrodes are named by their nominal design diameter. Values are mean *±* standard deviation over *n* = 3 (cauliflower) or *n* = 2 (pinecone) electrodes.

Impedance spectra are shown in Figure 2c and Figure 2d (magnitude of impedance and phase angle, respectively). As expected, the impedance varies with the size and type of the electrodes. The impedance magnitude at 1 kHz is approximately 3.28 kΩ and 15 kΩ, for 700 µm and 300 µm diameter “cauliflower” electrodes, respectively, and 1.8 kΩ and 6.1 kΩ for the corresponding “pinecone” electrodes, which falls within the typical range reported for gold thin-film electrodes of comparable size [55, 56]. The phase spectra indicate predominantly capacitive behavior at lower frequencies for all four electrode types.

Cyclic voltammograms of representative electrodes are shown in Figure 2e. The charge storage capacity (CSC) and its cathodic component (CSC_c_), together with the maximum cathodic charge injection capacity (CIC) obtained from voltage transient measurements (Figure S5, Supporting Information), are summarized in Figure 2f. Adding the “pinecone” structure increased both the stored and the injectable charge by approximately a factor of two at both diameters. For 700 µm electrodes, CSC_c_ increased from 4.99(129) mC cm^−2^ to 10.73(154) mC cm^−2^ and the CIC from 23.1(65) µC cm^−2^ to 49.9(30) µC cm^−2^; for 300 µm electrodes, CSC_c_ increased from 5.34(97) to 10.46(402) mC cm^−2^ and the CIC from 20.4(16) to 44.7(72) µC cm^−2^. Across the three measurements and both diameters the improvement ranged from 1.8 to 2.5 times, consistent with the increase in electrochemically active surface area evident in the SEM images. Values normalized per unit geometric area are not directly comparable between electrodes of different diameter, since the current distribution over a disc electrode is concentrated at its perimeter and the perimeter-to-area ratio of the 300 µm electrodes is approximately 2.5 times that of the 700 µm electrodes. Comparisons between surface types are therefore made only at matched diameter. Values are reported as mean *±* standard deviation over *n* = 3 electrodes for the cauliflower types and *n* = 2 for the pinecone types; the CIC of the 300 µm cauliflower electrodes is the mean of two electrodes. Electrodes are referred to throughout by their nominal design diameter. The laser ablation process reduces the actual exposed diameter, as shown for indicative electrodes in Table S3, so the reported area-normalized values are conservative.

### 2.5 Biocompatibility and biostability of polyurethane-bioelectronic foils

Polyurethane substrates were fabricated with and without embedded gold tracks using the two polyurethane materials, Platilon U 4201 AU (polyether TPU, Covestro) and Bionate 80A (medical grade, polycarbonate-urethane, dsm-firmenich), and subjected to *in vitro* cytotoxicity and *in vivo* implantation studies, as described below.

#### 2.5.1 Assessment of in vitro cytotoxicity of Bionate 80A and Platilon U 4201 AU on differentiated SH-SY5Y cells

Cytotoxicity was assessed on differentiated SH-SY5Y neuroblastoma cells according to ISO 10993 [57, 58]. In the indirect test, cell viability quantified by MTT and NRU assays was not significantly reduced relative to untreated cells for either material (Figure 3a-(i-ii)), remaining well above the 70% threshold below which a material is considered cytotoxic under ISO 10993-5. No morphological alterations were observed after exposure to the material extracts (Figure 3a-iii). In the direct contact test, relative cell viability quantified by MTT likewise remained above 70% for both materials (Figure 3a-(iv-v)). To confirm that the differentiated phenotype was maintained, mRNA expression of *MAP2* and *NEFH*, encoding microtubule-associated protein 2 and neurofilament heavy chain respectively, was quantified by RT-qPCR. No significant differences were found between cells in contact with either material and unexposed control cells (Figure 3a-vi). Neither Bionate 80A nor Platilon U 4201 AU was therefore cytotoxic to differentiated SH-SY5Y cells under the conditions tested.

**Figure 3:**
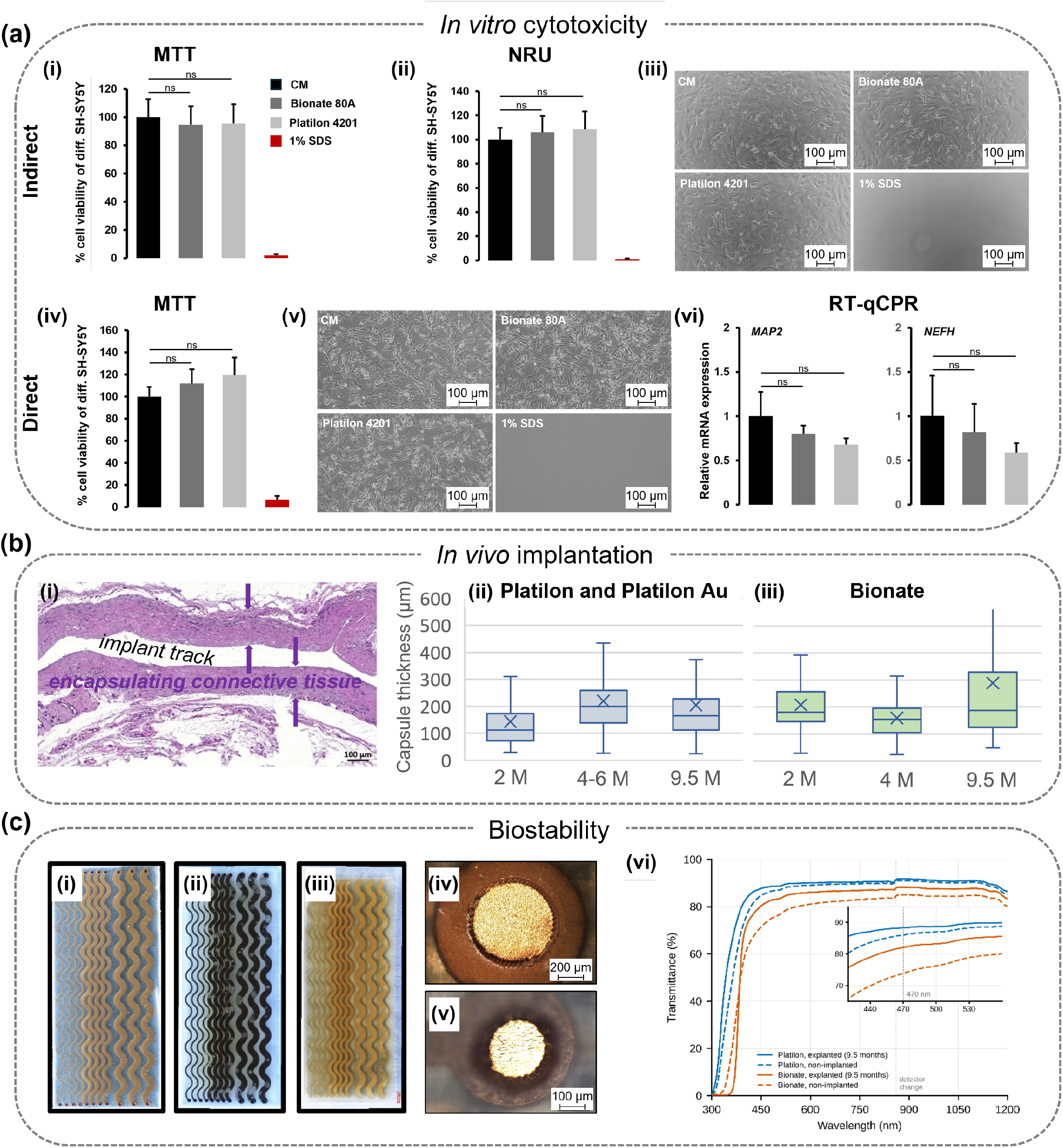
(a) In vitro cytotoxicity of Bionate 80A and Platilon U 4201 AU on differentiated SH-SY5Y cells. For indirect cytotoxicity, cell viability was quantified by the MTT (i) and NRU (ii) assays. Representative photographs of microscopic evaluation of differentiated SH-SY5Y following treatment with liquid extracts of materials for 24 hours (iii). Quantitative assessment of direct contact cytotoxicity test by MTT assay (iv). Representative photographs of differentiated SH-SY5Y upon direct contact with Bionate 80A or Platilon U 4201 AU for 40 hours are presented (v). RT-qPCR analysis of *MAP2* and *NEFH* mRNA expression. Relative gene expression levels were normalized to *HPRT1* (vi). Abbreviations: CM, culture medium; SDS, sodium dodecyl sulfate solution; ns, not significant. (b) Histological (H&E) image of the encapsulating connective tissue formed around the implanted sample (i). Thicknesses of the encapsulating connective tissue for pure Platilon TPU and Platilon TPU with gold electrodes (ii) and for Bionate (iii). The box plots also indicate the mean (cross sign), and the median (horizontal line inside the box). (c) Polyurethane sample with embedded gold electrodes before (i), and after 9.5 months of implantation (ii-iii), seen from the top (ii) and bottom (iii), together with the zoomed-in image of two electrodes/openings of different diameters (iv,v), showing no damage to the electrode after the implantation. Optical transmittance of the Platilon and Bionate TPU samples, before and after 9.5 months of implantation (vi).

#### 2.5.2 *In vivo* implantation

Nine samples were implanted subcutaneously in six female Wistar rats for periods between 2 and 9.5 months (Table S5). Three animals received pure TPU films bilaterally, with Bionate 80A on one side and Platilon U 4201 AU on the other, allowing the two materials to be compared within the same animal at each time point. The remaining three animals each received a single Platilon foil with embedded gold tracks and laser-ablated electrode openings. Because the gold-bearing foils carried open electrodes, exposed gold was in direct contact with the tissue. However, exposed gold accounted for less than 3% of the sample surface and gold is not expected to elicit a tissue response distinct from that of the surrounding polymer, the pure and gold-bearing Platilon groups were therefore grouped together.

The encapsulating connective tissue was visualized by haematoxylin and eosin (H&E) staining and its thickness measured every 500 µm along the track left by the explanted sample (Figure 3b-i); medians, interquartile ranges and the number of measurement points for each group are given in Table S6.

For Bionate 80A the tissue capsule thickness was 181 µm [147–252] at 2 months, 155 µm [106–195] at 4 months and 188 µm [126–318] at 9.5 months. For Platilon U 4201 AU the corresponding values were 111 µm [73–171] at 2 months, 199 µm [139–259] over a pooled 4–6 month window and 166 µm [113–228] at 9.5 months (Figure 3b-ii). Neither material showed a monotonic change with implantation time, interquartile ranges overlapped substantially across all time points, and with a single sample per material per time point no statistical comparison between materials or between time points was performed; the values are reported descriptively. The values measured at 9.5 months are of the same order of magnitude as those reported for other subcutaneous implants [59–61].

Histologically, the response was mild throughout. Eosinophils and neutrophils were only rarely visible, no tissue necrosis was observed in the vicinity of any implant, and at 9.5 months the capsule consisted of mature fibrotic tissue containing few fibroblasts.

#### 2.5.3 Biostability

Explanted foils were examined by optical microscopy (Figure 3c-(i–v)). No fracture or interruption of the gold structures was observed, although local delamination of the gold from the surrounding TPU was seen in places. Conductivity measurements after explantation confirmed that the tracks remained electrically continuous, with a resistance below 20 Ω. A similar local delamination was observed during soak testing of embedded tracks in PBS at 37 °C and 67 °C. In these tests gold delamination from the TPU developed over weeks to months, without interruption of the tracks (Figure S1; Supporting Information, Section SA.2).

The optical transparency of the TPU films was measured after 9.5 months of implantation and compared with non-implanted reference films of the same materials (Figure 3c-vi). Both materials transmit strongly across the visible and near-infrared range, with an absorption edge in the near ultraviolet. At 470 nm, a wavelength commonly used for optogenetic stimulation, the explanted films transmitted 88% (Platilon U 4201 AU) and 82% (Bionate 80A); the corresponding non-implanted reference films transmitted 86% and 74%. No reduction in transmittance was observed after implantation for either material. The reference films were separate coupons rather than the same films measured before implantation, so the small differences between the two conditions cannot be attributed to implantation. Both materials therefore retain high transparency after chronic implantation, supporting their use in neural interfaces for optogenetic-related applications, as well as in combination with optical imaging techniques.

#### 2.5.4 Removal of the sacrificial copper substrate

The process uses a Cu sheet as sacrificial substrate, which is removed by wet etching before the second TPU layer is laminated. Incomplete removal would leave Cu within a device intended for chronic implantation, so the elemental composition of finished foils was examined by energy-dispersive X-ray spectroscopy (EDX; Section 5). Four foils with embedded gold tracks and electrodes were analyzed: two non-implanted, one of which had been ultrasonicated for 15 minutes to remove organic surface contaminants (E1) and one of which was analyzed as received (E2), and two foils recovered after 5.5 and 9.5 months of implantation (E3 and E4).

In all four foils, copper was below the detection limit of the measurement, conventionally taken as approximately 0.1 wt% (Table S7). Applying this limit as an upper bound on the copper concentration and extending it to the whole polymer volume, the copper content of a single foil of the dimensions used here (mass 92.1 to 121.3 mg) does not exceed approximately 120 µg. Because no sample yielded Cu above the detection limit, the analysis does not distinguish between the ultrasonicated and non-ultrasonicated foils, or between the non-implanted and explanted foils. The wet etch therefore removes the sacrificial copper to below the sensitivity of this measurement.

### 2.6 Polyurethane-based bioelectronic foils with embedded silicon chips

Active bioelectronic foils were fabricated by embedding silicon chips within the TPU layers using the process described above (Figure 4). The process supports gold tracks 15 µm wide at a pitch of 30 µm (Figure 4a, inset), patterned without photomasks and covered by the merged polymer. Integrating a chip requires that the flip-chip bonding survives the subsequent lamination steps and that good contact is achieved between the chip pads and the substrate metallization. Dummy chips carrying daisy-chain and four-point measurement structures were used to assess this, allowing the bonded contacts to be characterized electrically after embedding (Figure S3a).

**Figure 4:**
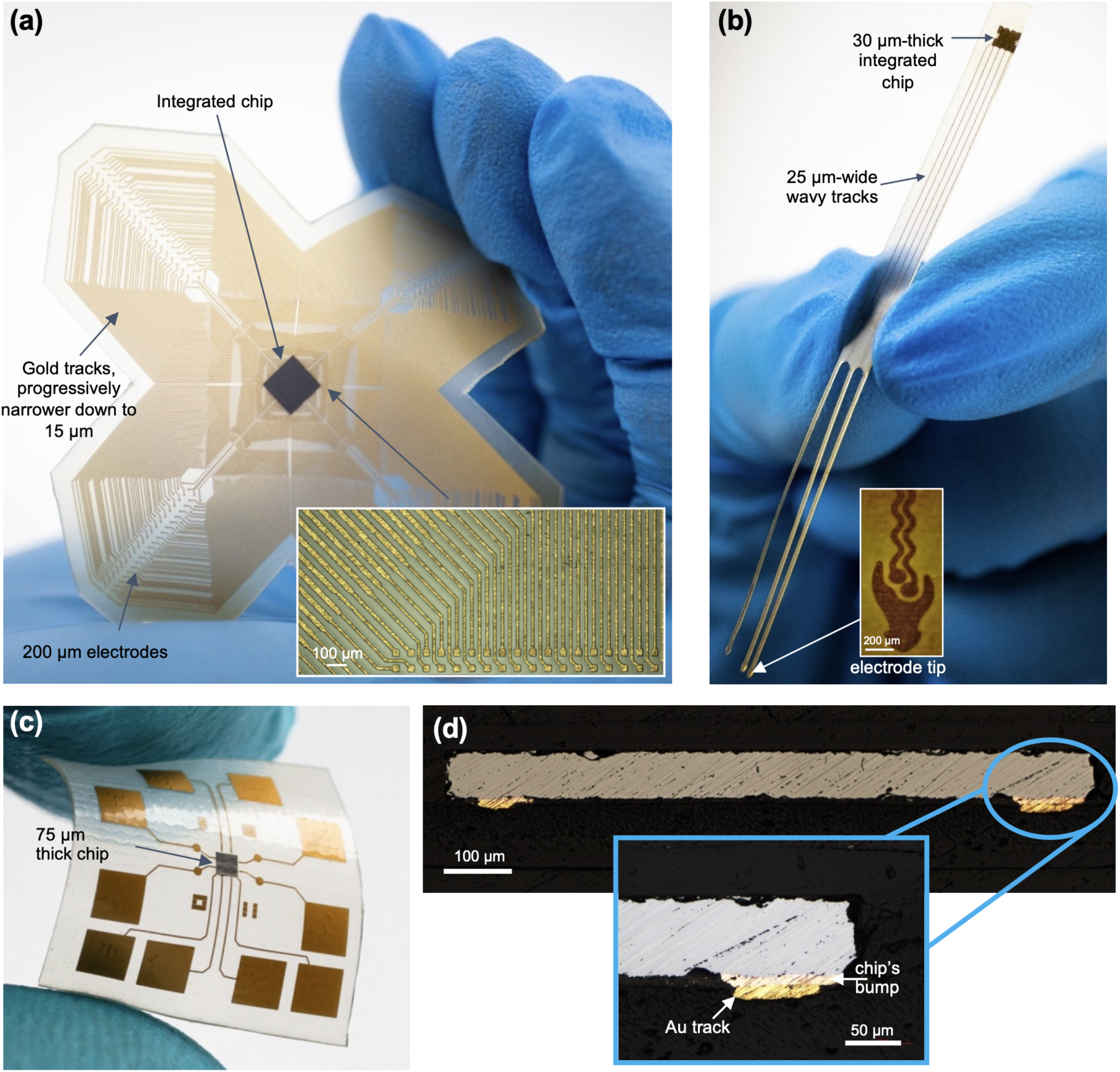
TPU-based bioelectronic foils with embedded silicon chips: (a) high-density foil intended for the central nervous system with an embedded 300 µm thick daisy-chain chip contacting 324 electrodes, (b) thinned (30 µm) dummy chip embedded in bioelectronic foil intended for the peripheral nervous system, (c) 950 µm × 950 µm, 75 µm thick daisy-chain chip in a substrate intended for electrical characterization, and (d) cross-section of its contacts (5 µm electroplated copper) connected to the substrate’s gold tracks.

Two underfill systems were used: an anisotropic conductive adhesive (ACA), and TPU applied as a non-conductive film (NCF). The latter was explored in order to minimize the number of materials and interfaces in the active prototypes. In this case, the volume beneath the chip is filled with the same material as the surrounding encapsulation, so that the assembly contains no adhesive and no additional material interface at the chip.

Chips of markedly different geometries were embedded, ranging from 4 mm × 4 mm × 300 µm (Figure 4a) to 950 µm × 950 µm × 75 µm (Figure 4c), together with a thinned 1.4 mm × 1.4 mm × 30 µm dummy chip (Figure 4b). The dies also carried a variety of pad metallizations, i.e., gold/palladium stud bumps (Figure S3b-i, inset), electroplated gold (Figure S3c-i, inset) and electroplated copper (Figure 4d, inset), further demonstrating the flexibility of the approach. An overview of the various chips used in this work and their characteristics can be found in Table S2.

The smallest die, a 950 µm × 950 µm × 75 µm daisy-chain chip with electroplated copper bumps (105 µm × 85 µm ×5 µm), was embedded using the more conventional ACA process (Figure 4c). A cross-section of a bonded contact after embedding is shown in Figure 4d.

To assess the TPU-as-NCF route, two custom daisy-chain chip layouts were assembled on TPU substrates carrying the complementary layout, at low and high contact density (Figure S3b,c). The low-density variant, chip “8072” (Figure S3b), carried 72 gold stud-bumped contacts on 80 µm pads on a 500 µm-thick silicon die, while the high-density variant, chip “361008” (Figure S3c), carried 1008 electroplated gold contacts of 36 µm × 36 µm at 80 µm pitch on a 300 µm-thick silicon die.

Four-point measurements on the low-density chip gave individual contact resistances between 4.5 and 14 mΩ, with all measured contacts conducting. On the same sample, a two-point measurement of the 64-contact daisy chain gave 18Ω, corresponding to an average contact resistance of approximately 0.25Ω once the estimated chip and substrate track contributions were subtracted. The two values differ by more than an order of magnitude, indicating that the individually addressable contacts characterized by the four-point method were not representative of every bond in the chain, consistent with the incomplete yield observed on the high-density chip.

For the high-density chip, contact resistances ranged from 5.5 to 73 mΩ, and two-point measurements across three assembled samples indicated a bond yield of approximately 90% for the outer ring of contacts, estimated from the fraction of measurement paths, i.e., each passing through two contacts, that failed to conduct (Figure S4). Further optimization of the bonding parameters will be required to improve this yield.

A demonstrator was assembled using the same high-density chip design, routing 324 of its contacts to peripheral electrodes through a single gold metallization layer, with the chip fully embedded in TPU (Figure 4a). This device was produced without electrode openings and its interconnections were therefore not characterized electrically; the resistance and yield data above were obtained on samples carrying the same chip bonded to substrates designed for four-point and daisy-chain measurement.

Different form factors can be addressed with the same approach. Figure 4b shows a probe intended for the peripheral nervous system, in which the interconnects are routed as 25 µm wide serpentine tracks and the thinned 30 µm dummy chip serves as a mechanical and geometrical placeholder. Depending on the functionality of the integrated chip, such foils could support multichannel recording or stimulation over large or multiple neuronal areas, or from small volumes at high resolution. The chips embedded in this work were daisy-chain and dummy structures without active circuitry, so the demonstration concerns the integration and interconnection process rather than device function.

## 3 Discussion

The process described here produces gold tracks of 15 µm width at 30 µm pitch on a soft thermoplastic substrate. This resolution is achieved because the metallization is patterned by photolithography on a rigid sacrificial substrate and only afterwards transferred into the polymer, rather than being deposited onto a compliant surface. Methods commonly used to metallize pliable substrates, such as inkjet printing, soft lithography or screen printing, do not reach comparable feature sizes. Conventional photolithography, however, is slow and costly, and requires a new photomask for each design. We therefore adapted laser direct imaging, established in printed circuit board manufacturing, in which the pattern is written directly into the photoresist from a digital layout. Combined with polymer processing, this yields a readily adjustable and inexpensive route to high-density metallization on a soft substrate.

Because TPU is a thermoplastic elastomer, successive layers merge under heat and pressure into a single continuous body rather than adhering across a boundary, as confirmed by the cross-sections in Figure 1d. This differs from the annealing used to improve parylene-to-parylene adhesion, which strengthens an interface that nonetheless remains present, and from self-bonding approaches based on dynamic covalent bond exchange, in which bonding is achieved through reversible chemistry within a crosslinked network [39]. Here the two layers become indistinguishable, and no interface remains at which delamination could initiate.

Electrochemically, the additional plating step used to produce the “pinecone” surface approximately doubled both the cathodic charge storage capacity and the charge injection capacity, at both electrode diameters and across all three measurements. For 700 µm electrodes, the cathodic charge storage capacity rose from 4.99 to 10.73 mC cm^*−*2^ and the charge injection capacity from 23.1 to 49.9 µC cm^*−*2^, with comparable gains at 300 µm. The improvement is obtained without altering the electrode footprint, which matters for high-density arrays where the available area per site is fixed.

Polyurethanes have several decades of clinical use in long-term implants. In pacemaker lead insulation using polyether-based grades, which are more vulnerable to oxidative attack, degradation modes such as environmental stress cracking, hydrolysis of the soft segment, and oxidation catalyzed by metal ions have been observed. This understanding has motivated the development of polycarbonate-urethanes. The two materials examined here span this distinction: Platilon U 4201 AU is a polyether TPU and Bionate 80A a medical-grade polycarbonate-urethane. Over 9.5 months of subcutaneous implantation, neither showed evidence of macrophage-mediated degradation, tissue necrosis was absent, and the optical transmittance of explanted films was not reduced relative to non-implanted references. These observations are consistent with both grades remaining chemically stable over this period, although a subcutaneous pocket is a less oxidatively demanding environment than direct contact with neural tissue, and 9.5 months is short relative to the intended service life of a chronic implant.

The element that did show change over time was not the polymer but the metal–polymer interface. Soak testing of embedded tracks in PBS produced local delamination of the gold from the surrounding TPU over weeks to months (Supporting Information, Section SA.2), and comparable local delamination was seen on explanted foils. In neither case were the tracks interrupted, and explanted foils remained electrically continuous, which is consistent with the surrounding merged polymer holding the metallization in place even where adhesion has been lost. Over longer periods, however, a delaminated interface offers a path for fluid ingress along the track and is a plausible route to eventual corrosion. Improving gold adhesion to TPU is therefore the most direct means of extending device lifetime. Because the process uses a sacrificial copper substrate, residual copper was assessed by elemental analysis. Copper remained below the detection limit of the technique in all samples, placing an upper bound of approximately 120 µg on the copper content of a single foil, which is more than an order of magnitude below the daily Adequate Intake proposed by the European Food Safety Authority [62]. The bound is conservative, since the analysis is confined to the first micrometre below the surface, where any copper remaining after etching would be concentrated. It nonetheless constrains the copper present rather than the fraction available for release, which is what determines local tissue exposure. The absence of necrosis and the mild inflammatory response observed histologically are more direct evidence of tissue exposure.

Thermoplastic encapsulation with liquid crystal polymer derives its performance from both a fusible interface and an intrinsically low water vapor transmission rate. Polyurethanes are more permeable to water in comparison, and the elimination of the polymer–polymer interface addresses delamination concerns rather than permeation. Silicon integrated circuits have been shown to be inherently hermetic, remaining electrically stable over a year of accelerated *in vitro* and *in vivo* exposure even as bare die [63], and polydimethylsiloxane, itself a moisture-permeable elastomer, limits chip degradation sufficiently to serve as a standalone encapsulant for years-long implantation provided its adhesion to the chip surface is maintained [63, 64]. Consistent with this, a comparison of thin-film encapsulants on LCP found that the water vapour transmission rate was not the main factor determining lifetime, and that adhesion to the substrate governed how long a stable interface was retained [32]. Where an additional barrier is required, thin inorganic layers could supply it. In our prior investigation, a 100 nm hafnium-based ALD multilayer showed no ionic penetration after seven months of direct exposure to the body environment [65], and such coatings are compatible with the temperatures used here.

The concern of the lower barrier properties for passive metallization relates to the electrical performance, rather than corrosion. Water uptake raises the permittivity of the dielectric and thus the capacitive coupling between neighbouring tracks. However, this effect is modest at realistic uptake levels expected. The more significant risk is ionic permeation, which lowers insulation resistance and introduces resistive leakage between conductors. At the 30 µm pitch demonstrated here the conductor spacing is small enough for this to potentially affect crosstalk between channels. A delaminated metal–polymer interface is of particular concern, since it offers a continuous path for fluid along the length of a track. We have previously shown that PDMS, although permeable to water vapour, effectively blocks the ionic species present in the body [63]. The corresponding behaviour of TPU has not, to our knowledge, been characterized. Measurement of insulation resistance and interchannel crosstalk over soak time is therefore a necessary step before chronic active implants can be built on this platform.

Several limitations bear on the scope of these results. The chips embedded in this work were daisy-chain and dummy structures without active circuitry, so the demonstration concerns integration and inter-connection rather than device function. The *in vivo* evaluation was subcutaneous and did not involve contact with neural tissue, and the *in vitro* work used a neuroblastoma cell line; neither establishes performance at a neural interface. One sample was implanted per material per time point, so no statistical comparison between materials or time points is possible and the histological data are reported descriptively. No sham or incision-only group was included, so the measured capsule thicknesses have no surgical-trauma baseline. Foils with gold tracks could not be produced reliably in the thicker Bionate stack and were therefore implanted only in Platilon, so chronic *in vivo* data for the medical-grade material exist only for the bare film. Finally, the electrochemical characterization was performed on two to three electrodes per condition.

Taken together, the cross-sectional, soak, *in vitro* and *in vivo* results support thermoplastic polyurethane as a substrate and encapsulation material for soft neural interfaces, and the process described here as one route to realizing them. The mechanical compliance of TPU is substantially closer to that of neural tissue than that of polyimide or parylene C, and at the film thicknesses used here the resulting bending stiffness is correspondingly low, which is expected to reduce micromotion at the tissue interface. Using TPU itself as the underfill removes the last adhesive interface within the assembly, but bonding onto a thermoplastic substrate imposes constraints absent when bonding onto a rigid carrier: the substrate softens at the bonding temperature, and the chip must be held under pressure until it has cooled below the melting point of the polymer. Under the conditions used here, approximately one bond in ten did not conduct, and further optimization of the bonding profile would be required to improve the yield of highdensity assemblies. Lastly, longer-term studies will be needed to establish how effectively TPU protects embedded silicon. Such implants could in future be powered wirelessly through an integrated antenna or ultrasound transducer, removing the need for surgical battery replacement.

## 4 Conclusion

This work introduces thermoplastic polyurethane as a substrate and encapsulation material for soft neural interfaces, and demonstrates one route to realizing them. TPU combines properties not otherwise found together in a single material: decades of clinical use in long-term implants, availability in medical grade, a stiffness well below that of polyimide and parylene C, compatibility with microfabrication, and, as a thermoplastic elastomer, the ability to merge with itself under heat and pressure so that no polymer–polymer interface remains within the finished device.

Using a polymer-last process in which gold is patterned on a sacrificial copper substrate by maskless laser direct imaging and transferred into TPU by vacuum lamination, we produced conformally covered tracks down to 15 µm at 30 µm pitch within a single, merged polymer body. An additional electroplating step producing a nanostructured electrode surface approximately doubled the charge storage and charge injection capacities at both electrode diameters, and silicon chips were flip-chip bonded and fully embedded using TPU itself as the underfill. Both materials examined were non-cytotoxic to differentiated SH-SY5Y cells, and after 9.5 months of subcutaneous implantation the tissue response remained mild, with optical transmittance undiminished relative to non-implanted films.

The limiting element over time was not the polymer but its interface with the metal: local delamination of gold from TPU appeared during soak testing and on explanted foils, without interrupting the tracks. Improving this adhesion, and characterizing the ionic barrier performance of the merged TPU body, are the next steps toward chronic active implants on this platform.

## 5 Experimental section

### Patterning of TPU

Micropatterns were fabricated on 25 µm- and 50 µm-thick TPU foils (Platilon U 4201 AU, Covestro), using a UV-based picosecond laser (Picodrill, Schmoll) with a 20 µm laser diameter and the following process parameters: 3.3 W and 4 W for 25 µm and 50 µm TPU foil thickness respectively, using 13 repetitions at 200 kHz laser frequency.

### Bionate 80A solution preparation and film fabrication

Bionate 80A pellets were dried in an oven at 80 °C for 4 h. After drying, the pellets were weighed and the required amount of DMAc was calculated to obtain a 20% (w/w) Bionate 80A solution. The dried Bionate pellets were subsequently added to the DMAc in a beaker. The beaker was placed on a hot plate equipped with a thermostat and covered with a Teflon sheet to minimize evaporation of DMAc. A mechanical mixer with a mixing blade (MINISTAR 40 digital, IKA, Germany) was used for mixing at 500 rpm. The solution was initially mixed for 30 min before the temperature was gradually increased to 65 °C.

The solution was maintained at 65 °C until the Bionate was fully dissolved and was then cooled to room temperature. If required, the solution concentration was recalculated after heating based on the measured masses, as the mass of the beaker and Bionate remained constant while the mass of DMAc decreased.

The Bionate solution was filtered by pressing it through a 1 *µ*m filter into a metal or polypropylene syringe. A maximum of 70 mL of solution was filtered due to the capacity of the filter. Before film deposition, the solution was centrifuged at 2888 rpm for 1 min (Centrifuge 5804 R, Eppendorf, Germany) to remove air bubbles from the solution.

The solution was subsequently weighed and poured into a 100 mm-diameter Petri dish to obtain a target dry film thickness of 100 *µ*m after evaporation of the DMAc. Based on a Bionate 80A density of 1.19 g cm^*−*3^, a Bionate mass of approximately 0.935 g was required to obtain a 100 *µ*m-thick film over the 100 mm-diameter area. For the 20% (w/w) Bionate solution, this corresponds to approximately 4.67 g of solution, consisting of 0.935 g Bionate and 3.74 g DMAc. The solution was then left in the Petri dish for evaporation of the DMAc.

### Polymer-last fabrication process

A 70 µm thick roughened copper (Cu) sheet with up to 2 µm roughness was used as a sacrificial substrate (Figure 1c-1), and was laminated on both sides with a 25 µm dry film photoresist (RD1225, Hitachi) (Figure 1c-2). Maskless lithography was used to pattern the resist (Figure 1c-3), leaving exposed the areas of Cu that would later form the tracks, electrodes and pads. A gold (Au) layer (5 - 20 µm thick) was deposited through an electroplating process on the exposed Cu (Figure 1c-4); the electroplating conditions and the resulting layer thickness are given in Section 5. The soft photoresist mask was then removed chemically (Figure 1c-5), in a mixture of 2-aminoethanol (C_2_H_7_NO) (2 %) and potassium hydroxide (KOH) (10 %) at 50 °C. At this point, a thin silicon chip was flip-chip bonded to the Au pads using an underfill or adhesive. Variations included: an ACA with 3 µm Au-coated polymer sphere particles (TOSHIBA, TAP0201C), or a Platilon TPU film of varying thickness used as a NCF underfill (Figure 1c-6). The bonding process was carried out on a SET FC 150 FAV flip-chip bonder with temperatures reaching 180 - 190 °C and bonding forces between 4000 and ∼7500 g, depending on the chip size and selected underfill (some representative bonding profiles can be found in Figure S3, Supporting Information). Next, the first TPU layer (Platilon U 4201 AU, Covestro or Bionate 80A, dsm-firmenich) was laminated on top, together with an FR4-Teflon-TPU support stack (Figure 1c-7). Lamination was carried out in vacuum at 180 °C and 100 N cm^*−*2^; the full lamination profile is shown in Supporting Information, Figure S2a. The Cu sacrificial layer was subsequently etched in a copper chloride (CuCl_2_), hydrogen chloride (HCl) and hydrogen peroxide (H_2_O_2_) solution at 50 °C, and then rinsed with deionized water (Figure 1c-8). The stack was then flipped and a second TPU layer was laminated onto the exposed side to electrically insulate the Au structures (Figure 1c-9), at 165 °C and 100 N cm^*−*2^; the profile is given in Figure S2c of the Supporting Information). Openings for the electrodes and pads were produced with a picosecond laser (Picodrill, Schmoll) (Figure 1c-10), having a frequency range from 200 kHz up to 1 MHz, in two steps: an initial ablation, performed with the focal plane at the height of the substrate surface (1 W, 200 kHz, 7 repetitions) followed by a post-processing cleaning step with the focal plane raised by 0.4 mm (1 W, 400 kHz, 2 repetitions). Further cleaning steps with the same parameters were applied after raising the focal plane in 0.1 mm increments, until no TPU particles remained visible on the gold electrode surface. The heat generated during ablation causes the TPU to melt and reflow at the edge of the opening, such that the exposed electrode diameter is smaller than the nominal diameter of the laser pattern [40]. Nominal and measured diameters for some indicative electrodes are compared in Table S3 of the Supporting Information. An additional electroplating step could be applied at this stage to microstructure the electrode surface and increase its effective surface area, producing the “pinecone” electrodes described in Section 5. Finally, the sample was released from the support board by laser-cutting the TPU (Figure 1c-11). The teflon foil present between the FR4 board and the TPU substrate facilitated the release of the samples. Passive electrode arrays were fabricated by omitting the chip-bonding step.

### Gold metallization and electrode surface structuring

All Au tracks, electrodes and pads were electroplated in an electrolytic cell using a platinised titanium grid as the anode and the sample itself as cathode. The plating was performed galvanostatically with a source meter (Keithley 2400, Keithley Instruments) in a conventional soft Au electrolyte (Puramet 202, Schloetter) at 60 °C, pH 6.8 and 2.5 mA cm^*−*2^ for 33 min, yielding a 5 µm thick Au layer. A constant convective heat transfer was maintained through stirring of the solution. Layer thicknesses between 5 and 20 µm are accessible by varying the plating time. Two types of electrode surfaces were produced. “Cauliflower” electrodes (Figure 2a) were obtained directly after laser ablation of the electrode openings, without further processing. “Pinecone” electrodes (Figure 2b) were obtained by applying an additional electroplating step on 3.2 µm “cauliflower” electrodes after laser ablation, using gravitation-assisted plating without external convection, in the same electrolyte solution (Puramet 202, Schloetter), at 50 °C, pH 7.5, 2.5 mA cm^*−*2^, 11 min). This step is described in more detail in [66]. The resulting 1 µm thick nanostructured layer increased the electrochemically active surface area of the electrode.

### Electrode characterization

Electrical characterisation was performed on passive structures, i.e. Au tracks embedded in TPU without a bonded chip, fabricated as described in Section 5 with 700 µm and 300 µm nominal openings. Electrodes are referred to by their nominal design diameter throughout. Area-normalized quantities were calculated using the nominal electrode diameters, since measured opening diameters were available only for a subset of electrodes. Electrical contact between the samples and the potentiostat was established by inserting the interconnection pads into a ZIF connector (52271, Molex), whose soldered wires were connected to a Modulab XM potentiostat (Solartron Analytical, AMETEK). Measurements were carried out in a three-electrode configuration with a silver/silver chloride (Ag/AgCl) reference electrode (RE) and a 2 cm ×2 cm × 0.015 cm platinum (Pt) sheet counter electrode (CE). All electrodes were immersed in phosphate-buffered solution (PBS) (pH 7.4, Carl Roth) at room temperature. Grounding was set internally through the potentiostat. Electrochemical impedance spectroscopy (EIS) was performed between 1 Hz and 100 kHz with an excitation voltage of 10 mV.

Cyclic voltammograms were recorded in the same three-electrode configuration and electrolyte as the impedance measurements, at a scan rate of 50 mV s^*−*1^ within the water window of Au, between −0.8V and 0.6 V vs. Ag/AgCl. Fifty cycles were recorded to allow the electrode to reach a steady state, and the second-to-last cycle was used for analysis. The charge storage capacity (CSC) was obtained from the time integral of the current over one cycle, normalised to the electrode surface area; for the cathodic charge storage capacity (CSC_c_), only the negative (cathodic) portion of the curve was integrated.

Voltage transient measurements were performed to estimate the charge injection capacity (CIC). An asymmetric, cathodic-first, charge-balanced biphasic current pulse with a 1:4 amplitude ratio was applied between the working and counter electrodes, with a cathodic phase width of 200 µs and an inter-phase delay of 20 µs. The current waveform was generated by a custom-built voltage-controlled current source, driven by the arbitrary function generator of a mixed-domain oscilloscope (MDO34, Tektronix); voltage transients were recorded between the working and reference electrodes on the same instrument. The current amplitude was increased stepwise until the polarisation across the electrode interface reached the cathodic limit of the water window (− 0.8 V vs. Ag/AgCl). The maximum cathodic CIC was calculated as the product of the maximum current amplitude and the cathodic phase width, divided by the electrode surface area.

### Cell culture and SH-SY5Y differentiation

SH-SY5Y neuroblastoma cells, provided as a gift by Dr. Kostas Vekrellis (Biomedical Research Foundation, Academy of Athens, Athens, Greece) [67], were grown in Dulbecco’s Modified Eagle Medium (DMEM, Gibco) supplemented with 10% fetal bovine serum (FBS, Gibco), 1% penicillin/streptomycin (Gibco), and differentiated in the presence of 10 *µ*M retinoic acid (RA) for 7 days in a humidified incubator at 37 °C, in an atmosphere of 5% CO_2_, as described [68]. Differentiation was confirmed morphologically and by quantification of the relative expression levels of the neuronal markers *MAP2* and *NEFH* (Section 5).

### Indirect cytotoxicity test

Material liquid extracts were prepared in accordance with ISO 10993-12 [58]. Specifically, the materials were immersed in serum-free culture medium for 72 hours at 37 °C and 5% CO_2_ at a ratio of 6 cm^2^ of material per 1 mL of medium, after which the extracts were collected and supplemented with 10% FBS. SH-SY5Y cells were seeded at a density of 1 *×* 10^4^ cells per well in 96-well plates and differentiated as described in Section 5. The conditioned media were then replaced with the extracts and the cells were incubated for 24 hours at 37 °C in 5% CO_2_. Cytotoxicity was assessed qualitatively under a phase-contrast microscope (Carl Zeiss) and quantified by both the 3-(4,5-Dimethylthiazol-2-yl)-2,5-diphenyltetrazolium bromide (MTT) and neutral red uptake (NRU) assays, according to ISO 10993-5 [57].

### Direct contact cytotoxicity test

The direct contact cytotoxicity test was conducted in accordance with ISO 10993-5 [57]. Shortly, SH-SY5Y cells were cultured and differentiated on 6-well plates (approximately 10 cm^2^ surface area per well). A 1 cm^2^ piece of the tested material was placed on the center of each well, followed by incubation in the presence of the material for 40 hours at 37 °C and in 5% CO_2_. Then, the cells were observed under the microscope for morphological alterations, and cell viability was quantified with MTT, as described above.

### Reverse Transcription Quantitative PCR (RT-qPCR)

Total RNA were extracted from differentiated SH-SY5Y cells using the Nucleospin RNA kit (Macherey-Nagel) according to the manufacturer’s instructions. For cDNA synthesis, RNAs were reversely transcribed using SuperScript First-Strand Synthesis System (Invitrogen), and the resulting cDNAs (25 ng) were amplified using SYBR Select Master Mix (Applied Biosystems) and gene-specific primers targeting the Microtubule-associated protein 2 (*MAP2*) and Neurofilament heavy chain (*NEFH*) genes. Gene expression was normalized against the housekeeping gene *HPRT1* encoding for Hypoxanthine phosphoribosyltransferase 1. The gene-specific primer sequences are listed in Table S4, Supporting Information.

### In vitro statistics

Statistical results are expressed as median ± SD from 3 independent experiments. All reactions were performed in triplicate in each experiment. Statistical analysis was performed using GraphPad (10.4). The differences between groups were assessed using an unpaired t-test and p-value higher than 0.05 was considered not statistically significant.

### In vivo implantation and histology

*In vivo* implantation was performed to evaluate the biostability and biocompatibility of the fabricated samples and their suitability for long-term use. Because Platilon U 4201 AU (Covestro) is not certified as a medical grade material, samples prepared from medical-grade Bionate 80A (dsm-firmenich) were included alongside pure Platilon films and Platilon foils with embedded gold tracks (Figure 3c-i). In the gold-bearing foils the electrode openings were produced by laser ablation as described in Section 5, so gold was exposed at the electrode sites. All samples measured 20 mm ×8 mm and were laminated from five 100µm TPU sheets, giving a final thickness of approximately 400µm after compression during lamination. Prior to implantation, samples were submerged in over 98% 2-Propanol for at least 5 minutes, then thoroughly washed with sterile distilled water and immediately implanted into the prepared animal. Nine samples were implanted in six female Wistar rats (14 weeks old, obtained from Toxi-Coop, Budapest, Hungary), as detailed in Table S5. Three animals received two samples each, implanted bilaterally with Bionate 80A on the left side and Platilon U 4201 AU on the right; the remaining three animals each received a single Platilon foil with gold tracks. Side allocation was fixed rather than randomized. All procedures were approved by the Institutional Animal Care and Use Committee of the Research Centre for Natural Sciences and by the Hungarian National Scientific Ethical Committee on Animal Experimentation (permit number: PE/EA/1253-8/2019).

Rats were anesthetized with isoflurane gas (2–2.5%), and the samples were implanted subcutaneously through a dorsal incision. Implantation was uneventful in all animals, with slightly larger wounds required for two of the samples (Table S5). Animals were maintained for the survival times given in Table S5; the longest survival group was extended beyond the originally planned six months to 9.5 months. At the end of each survival period animals were euthanized by isoflurane overdose (5% isoflurane mixed in 100% oxygen in a sealed chamber until no breathing and heart beats present), the implants were retrieved and the surrounding tissue was excised. Tissue was immersion-fixed in a 4% paraformaldehyde solution, embedded in paraffin, sectioned and stained with haematoxylin and eosin (H&E) and trichrome for histological analysis. Because the implant was removed before processing, the capsule was identified as the connective tissue lining the residual void left by the sample. Its thickness was measured on H&E sections every 500µm along this track. With one sample per material per time point, no inferential statistical comparison between materials or between time points was performed. As our data did not follow a normal distribution (verified with the Kolmogorov–Smirnov and Lilliefors tests), results are reported as the median [first to third quartile] of the measurement points, together with the number of points per group (Table S6). For Platilon, the 4–6 month and 9.5 month groups combine pure films with gold-bearing foils, as detailed in Table S6.

### Elemental analysis

The elemental composition of the finished foils was determined by energy-dispersive X-ray spectroscopy performed in a field-emission scanning electron microscope (FE-SEM, Supra 55 VP, Zeiss, with an EDS X Max 80, Oxford) at accelerating voltages of 5 and 10 kV and a working distance of 8.5 mm. Six measurements were acquired per sample and averaged. One non-implanted foil was ultrasonicated in deionized (DI) water for 15 minutes prior to analysis in order to remove organic surface contaminants; a second non-implanted foil was analyzed as received. The two remaining foils were analyzed after explantation and removal of adherent tissue (Section 5).

### Optical transmittance

Optical transmittance was measured on pure TPU films using a UV/Vis/NIR spectrophotometer (Lambda 950, PerkinElmer, Waltham, MA, USA) over the range 300 to 1200 nm at 5 nm intervals. Explanted films (samples 9 and 10, Table S5, both recovered after 9.5 months) were compared with non-implanted reference films of the same two materials.

### Flip-chip assembly and electrical characterization

Dummy silicon chips were fabricated at the Else Kooi Laboratory (Delft University of Technology) [69]. Contacts were either stud bumped manually or electroplated at wafer level by an external supplier. Four chip designs were used: chip “8072”, 500µm-thick, with 72 AuPd stud-bumped contacts on 80µm pads; chip “361008”, 300µm-thick, 4 mm ×4 mm in area, with 1008 electroplated gold contacts of 36µm ×36µm at 80µm pitch and 50µm wide bumps; chip “DC-75”, a 950µm ×950µm× 75µm daisy-chain chip with 105µm ×85µm× 5µm electroplated copper bumps, and chip “RFID-30”, 1.4 mm ×1.4 mm and 30µm thickness used as dummy mechanical and geometrical placeholders. Full details of the chip variants are summarized in Table S2.

Chips were flip-chip thermocompression bonded to the gold pads using either an ACA containing 3µm gold-coated polymer spheres (TAP0201C, Toshiba) or a Platilon U 4201 AU TPU film as an NCF (Figure 1c-vi). NCF pieces were cut slightly larger than the chip footprint from sheets of 50µm and 25µm thickness for chips “8072” and “361008” respectively, cleaned in isopropanol and dried at 80 °C for 30 minutes to prevent void formation. Representative bonding profiles when TPU was used as a NCF underfill are reported in Figure S3, Supporting Information. After bonding, samples were laminated and the sacrificial copper etched as described in Section 5.

Contact resistance on chips “8072” and “361008” was measured in a four-point configuration, with current driven between one pair of pads and voltage sensed across a second pair connected to the same bond. Each contact can be addressed in four equivalent configurations, and measurements were repeated across these to exclude geometric artifacts. For chip “8072”, a daisy chain running through 64 of its 72 contacts, was additionally measured in a two-point configuration; the average contact resistance was obtained by subtracting the estimated resistances of the chip and substrate tracks from the total chain resistance and dividing by the number of contacts, with the substrate track contribution corrected for the porosity of the electroplated gold. For chip “361008”, bond yield was estimated from two-point measurements across the outer contact ring, each path passing through two bonds, by assuming equal failure probability for the two contacts in each path. The 950µm daisy-chain chip was assembled to demonstrate embedding at reduced footprint, while its bonds were assessed by cross-sectioning rather than electrically.

## Supporting Information

Supporting Information is available as a separate file on BioRxiv.

## Acknowledgements

The authors would like to thank Michael Gerberich, Andreas Grünschneder, Michael Zwanzig, and Punal Surani, for their support during numerous fabrication steps. AI assistants were used during the preparation of this manuscript for coding, editing and language polishing. The authors reviewed and are responsible for all content, code and experiments.

