## Supporting Information for "Thermoplastic polyurethane as a base material for soft, high-density neural interfaces"

A. Pak, A. I. Velea, T. Hosman, R. Panskus, P. de Anta Dardagan, N. Bakhshaee Babaroud, V. Giagka
Department of Microelectronics, Delft University of Technology, Delft, The Netherlands
A. Pak, A. I. Velea, T. Hosman, J. Wilson, R. Panskus, M. Seckel, P. de Anta Dardagan, R. Schmidt, A. Gollhardt, B. Pahl, C. Kallmayer, V. Giagka
Fraunhofer Institute for Reliability and Microintegration IZM, Berlin, Germany
D. Horváth, K. Tóth, D. Meszéna, I. Ulbert
HUN-REN Research Centre for Natural Sciences, Institute of Cognitive Neuroscience and Psychology, Budapest, Hungary
D. Horváth, D. Meszéna, I. Ulbert
Pázmány Péter Catholic University, Faculty of Information Technology and Bionics, Budapest, Hungary
E. Zingkou, A. Kolianou, G. Sotiropoulou
Department of Pharmacy, School of Health Sciences, University of Patras, 265 04 Rion-Patras, Greece
S. Negi, F. Solzbacher
Department of Electrical and Computer Engineering, University of Utah, Salt Lake City, UT 84112, USA
S. Negi, F. Solzbacher
Blackrock Neurotech, Salt Lake City, UT 84108, USA
F. Solzbacher
Department of Biomedical Engineering, University of Utah, Salt Lake City, UT 84112, USA
Department of Materials Science and Engineering, University of Utah, Salt Lake City, UT 84112, USA

### S1 Sample configurations

Various sample configurations were used in this work; they are summarized in Table S1 and are referred to by these designations throughout.

Table S1: Sample configurations used in this work.

| **Used in** | **Configuration** | **Material** | **TPU stack** | **Metallization** | **Notes** |
|---|---|---|---|---|---|
| §2.1 Softness and conformability | Laser-micropatterned film | Platilon | Single film, 25 and 50 µm | None | Micropatterned by picosecond laser |
| §2.3 Fusion and conformal coverage | Laminated stack, with and without tracks | Platilon | $2 \times 100$ µm; $5 \times 100$ µm | None, or with Au tracks | Non-merged control laminated at 145 °C; merged at 165 °C. Soaked 100 d in PBS at 37 °C, or implanted 6 months (sample 8, Table S4) |
| §2.4 Electrode characterization | Passive array | Platilon | $2 \times 100$ µm | Au tracks, 5 µm; cauliflower and pinecone surfaces | Nominal openings 700 and 300 µm |
| §2.5.1 *In vitro* cytotoxicity | Bare film | Bionate; Platilon | Single films | None | 1 $cm^2$ pieces for direct contact; extracts at 6 $cm^2$ $mL^{-1}$ |
| §2.5.2 *In vivo* implantation | Bare film | Bionate; Platilon | $5 \times 100$ µm | None | samples 1, 2, 5, 6, 9, 10 (Table S4) |
| | Foil with electrodes | Platilon | $5 \times 100$ µm | Au serpentines, 5 µm; 22 openings | 3 serpentines at 1000 µm, 4 at 500 µm, 4 at 50 µm, each with two open ends; samples 4, 8, 12 (Table S4) |
| §2.5.4 Elemental analysis | Foil with electrodes | Platilon | As above | As above | Mass 92.1–121.3 mg; two non-implanted (one ultrasonicated), two explanted |
| §2.5.3 Biostability | Explanted foil with electrodes | Platilon | As above | As above | Samples 4, 8, 12 (Table S4) |
| | Bare film, transmittance | Bionate; Platilon | Single films | None | Samples 9 and 10 (Table S4), explanted at 9.5 months, and non-implanted references |
| §2.6 Embedded silicon chips | Active prototype, CNS | Platilon | $6 \times 100$ µm | Au tracks to 15 µm at 30 µm pitch; 200 µm electrodes | "361008"-chip; TPU NCF underfill; no openings |
| | Active prototype, PNS | Platilon | $6 \times 100$ µm | 25 µm Au serpentine tracks | "RFID-30"-chip, ACA underfill; no openings |
| | Daisy-chain prototype | Platilon | $6 \times 100$ µm | Au 5 µm thick | "DC-75"-chip; ACA underfill |
| | Assembly test vehicles | Platilon | $8 \times 100$ µm | Complementary daisy-chain and four-point layout | "8072" or "361008"-chips; TPU NCF underfill, 50 µm and 25 µm sheets |
| §S2 Soak testing (SI) | Passive test structure | Platilon | $2 \times 100$ µm | Au tracks, 5 µm thick, 40 mm long | Widths 1000, 500, 200 and 50 µm; with openings |

Table S2: Characteristics of the silicon chips embedded in the polyurethane foils. All chips were dummy or test structures. The daisy-chain chips carry interconnected pads allowing the bonded joints to be tested electrically, whereas the RFID chip served as mechanical and geometrical placeholder.

| Chip | Type | Size (mm × mm × µm) | Contacts | Contact size (µm) | Pitch (µm) | Pad metallization |
|---|---|---|---|---|---|---|
| "8072" | daisy chain | 2.72 × 2.72 × 500 | 72 | 80 ⌀ | 80 | AuPd stud bumps |
| "361008" | daisy chain | 4 × 4 × 300 | 1008 | 36 × 36 (50 × 50 after electroplating) | 80 | electroplated Au, 10 µm |
| "DC-75" | daisy chain | 0.95 × 0.95 × 75 | 28 | 105 × 85 | 100 | electroplated Cu, 5 µm |
| "RFID-30" | dummy | 1.4 × 1.4 × 30 | 4 | 138 ⌀ | - | electroplated Cu with Sn finish, 15 µm |

#### S2 Soak testing

The adhesion of the gold tracks to the surrounding TPU was assessed by soaking embedded test structures in PBS at 37 °C and at 67 °C and inspecting them periodically by optical microscopy.
Test structures consisting of gold tracks embedded between two TPU layers were fabricated as described in Section **??**, following the route without chip bonding. The tracks were 5 µm thick, 40 mm long, and 1000, 500, 200 and 50 µm wide. Samples were submerged in PBS (pH 7.4, Carl Roth) and held in a humidity chamber at 37 °C or 67 °C. Top and bottom views were recorded by optical microscopy at intervals of at most 24 days for the samples held at 37 °C and 7 days for those at 67 °C. Total soak durations were 130 days at 37 °C and 100 days at 67 °C.
At 37 °C, tracks of 1000 and 500 µm width remained free of delamination for 76 days, with the first delamination observed at the following inspection on day 100; 50 µm tracks remained intact for 27 days, with delamination first seen on day 37. At 67 °C, 1000 µm tracks were intact for at least 37 days, with first delamination seen on day 48, and 50 µm tracks after 55 days, whereas 500 and 200 µm tracks showed no delamination for the full 98 days of observation. Delamination was local and did not interrupt the tracks, which remained electrically continuous throughout. Taken together with the local gold–TPU delamination seen on the explanted foils, these results identify the metal–polymer interface, rather than the polymer itself, as the element limiting the long-term integrity of the structures. The results do not support a quantitative extrapolation of lifetime. Delamination did not occur consistently earlier at the higher temperature, and did not depend monotonically on track width, so no acceleration factor can be derived from these data. Sample-to-sample variability is the most likely factor affecting these results.

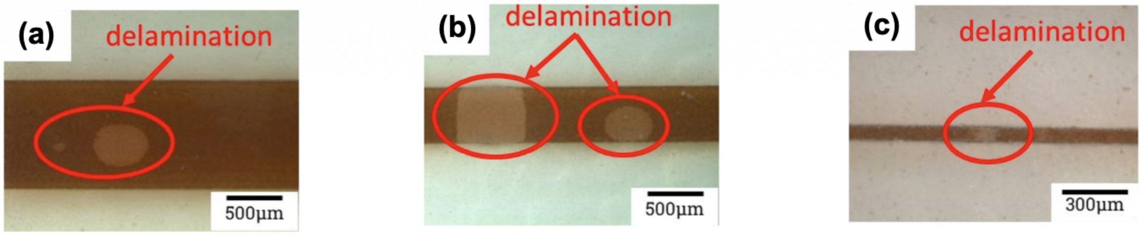


Figure S1: (a-c) Examples of the first delamination of gold tracks embedded in TPU after soak tests at 37 °C, started at day 100 (a), 100 (b), 37 (c) for line widths 1000, 500, 50 µm, respectively.

#### S3 TPU Lamination profiles

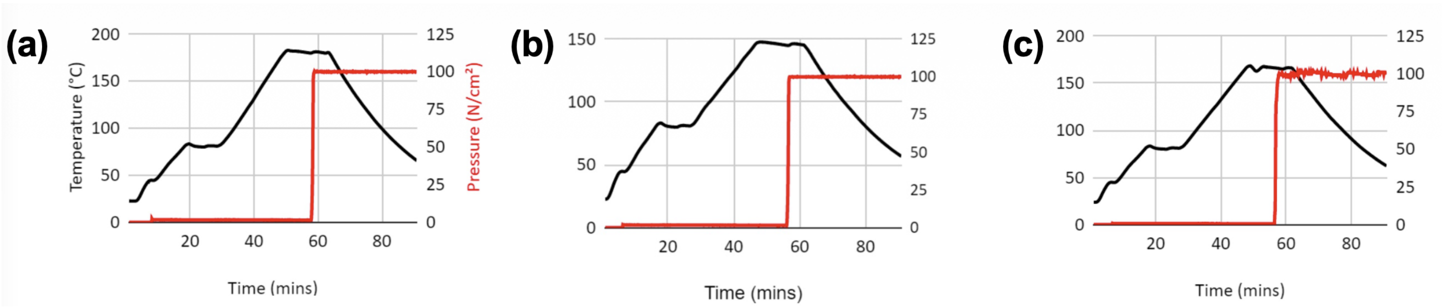


Figure S2: TPU lamination bonding profiles. (a) for the 1st lamination step (Figure 1c-7) used to bond the patterned gold electroplated on copper substrate to the TPU at 180 °C, (b-c) for the 2nd lamination step (Figure 1c-7) used to to cover exposed gold structures: profile for two non-merged TPU sheets laminated at 145 °C, shown in Figure 1d-i, 1d-v (b), and optimized profile for the merged TPU layers at 165 °C shown in Figure 1d-iii-iv, 1d-vi-viii (c). Lamination via thermocompression bonding was performed in the lamination press (MP30-VK-S, HML Haseneder Maschinenbau e.K.).

### S4 Bonding

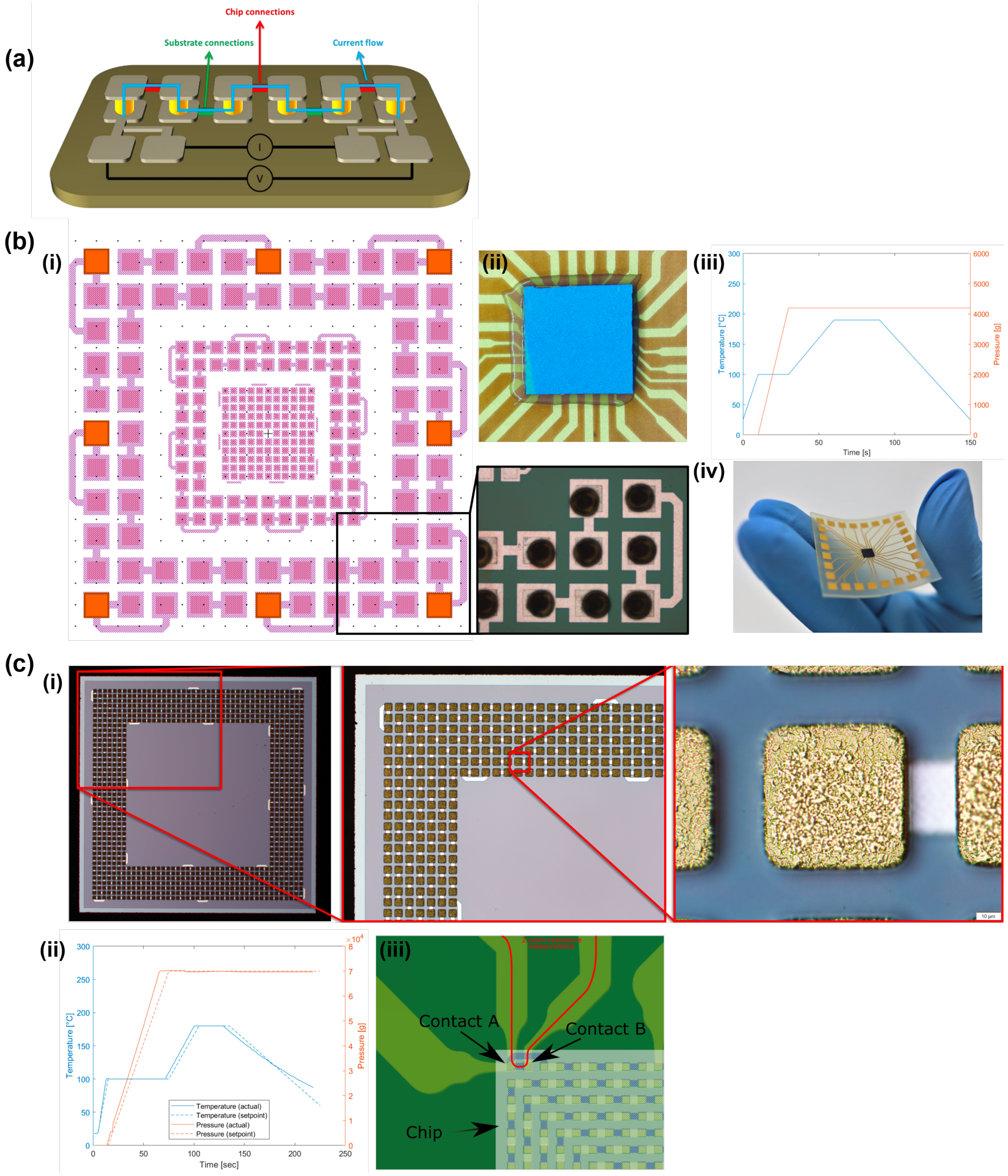


Figure S3: (a) Daisy-chain topology together with a four-point evaluation of it. (b)(i) Design of the low-density "8072" daisy-chain chip (only outer ring was used in this work, pad dimensions and pitch is 80 µm). Light-pink marks internal tracks of the chip, dark-pink marks exposed connections (pads), orange squares mark all pads on the outer ring that can be probed with a 4-point measurement. Zoom-in photograph shows part of the bumped ring. (ii) Top-view microscope image of a 500 µm flip-chip bonded "8072" chip with TPU as an NCF. (iii) Bonding profile used for the chip assembly through the TPU underfill. (iv) Photograph of the fully embedded chip in TPU. (c) (i) Photographs of the high-density daisy-chain "361008" chip with electroplated bumps. (ii) Bonding profile used for the chip assembly through the TPU underfill. (iii) Zoom-in of "361008" substrate render including chip (outline in white). The red line shows a conductivity line of a 2-point measurement. "Contact A" and "Contact B" indicate the two bonds through which the measurement runs and can be used to estimate bond yield.

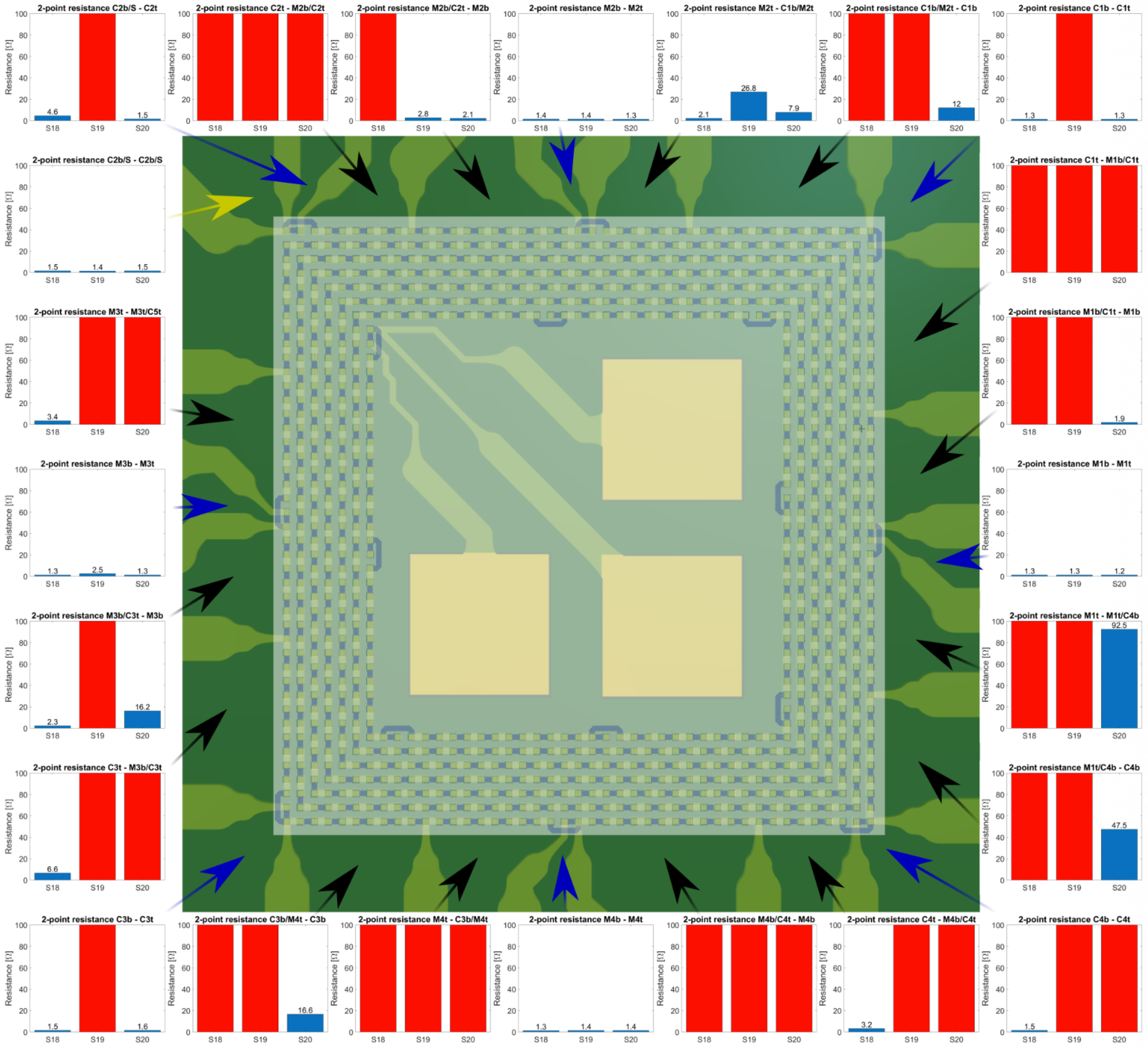


Figure S4: "361008" 2-point resistance measurements overview. In the center, a zoom-in of the substrate render (green and yellow colored) is shown with the chip configuration overlapping. The chip outline is shown transparent white, while its connections are shown in dark-blue. The outer picture area holds 2-point resistance measurement data of 3 samples. An arrow, originating from each of the graphs, marks between which two lines the measurement was conducted (the lines left and right of each arrow). Arrows in yellow indicate resistance measurements that do not run through the chip. Arrows in blue indicate 2-point measurements that run through the chip once. Black arrows indicate 2-point measurements that run through the chip multiple times (between 4-6 times). All resistances above 100 Ω are considered open-circuits and marked with red bars in the graphs.

#### S5 Electrical Characterization

During the TPU laser ablation to expose the gold electrodes, due to the heat produced by the laser, the TPU located on the borders of the opened circumference would melt and reflow, decreasing the actual exposed electrode diameter. To quantify this, in some indicative cases, the actual electrode diameter was calculated using optical microscopy images of the openings and compared to the diameter set on the laser. Results of the measurements are presented below:

Table S3: Nominal and measured electrode opening diameters. During laser ablation the TPU at the edge of the opening melts and reflows, reducing the exposed electrode diameter relative to the diameter set on the laser. Measured diameters were determined from optical micrographs of the openings for some indicative electrodes.

| Electrode | Nominal diameter (µm) | Measured diameter (µm) |
|---|---|---|
| Cauliflower, 700 µm | 700 | 620 |
| Cauliflower, 300 µm | 300 | 250 |
| Pinecone, 700 µm | 700 | 653 |

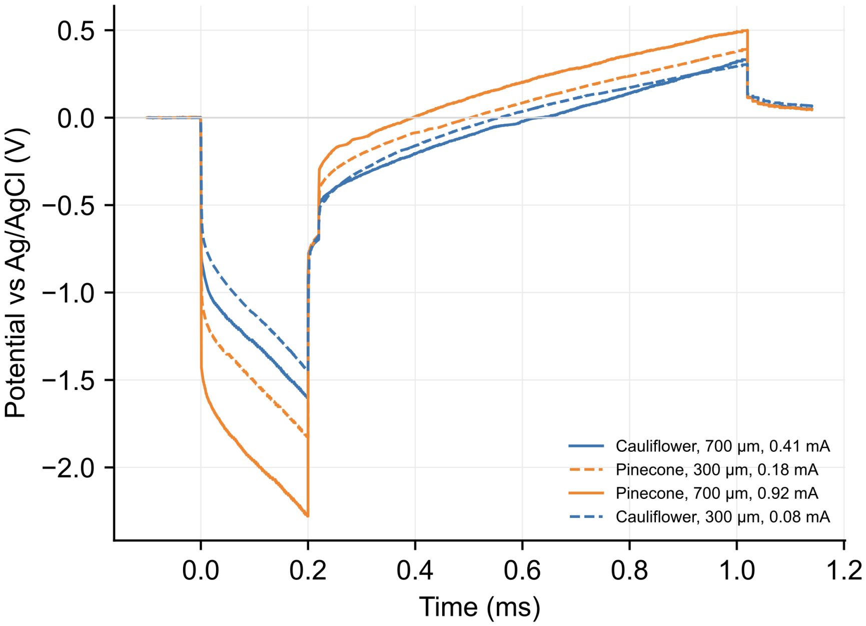


Figure S5: Voltage transient measurements for the 700 µm "cauliflower", 300 µm "cauliflower", 700 µm "pinecone" and 300 µm "pinecone" electrodes. Each electrode was pulsed at its own maximum current, as indicated at the legend.

### S6 Gene-specific primer sequences for RT-qPCR

Table S4: Sequences of the human gene-specific primers used for RT-qPCR. F = forward, R = reverse. *HPRT1* was used as the housekeeping gene for normalization.

| **Gene** | **Encoded protein** | **Primer sequence (5′ →3′)** |
| --- | --- | --- |
| *MAP2* | Microtubule-associated protein 2 | F: TCTGCCTCCTTCTCCACCCC<br>R: TCTGACTCCTTTTCCTTCTG |
| *NEFH* | Neurofilament heavy chain | F: CTGGAGGCACTGAAAAGCA<br>R: TCTTGACATTGAGCAGGTC |
| *HPRT1* | Hypoxanthine phosphoribosyltransferase 1 | F: GCCCTGGCGTCGTGATTAGT<br>R: AGCAAGACGTTCAGTCCTGTC |

### S7 Details of the *in vivo* implantation study aimed to evaluate the biocompatibility of the fabricated bioelectronic foils

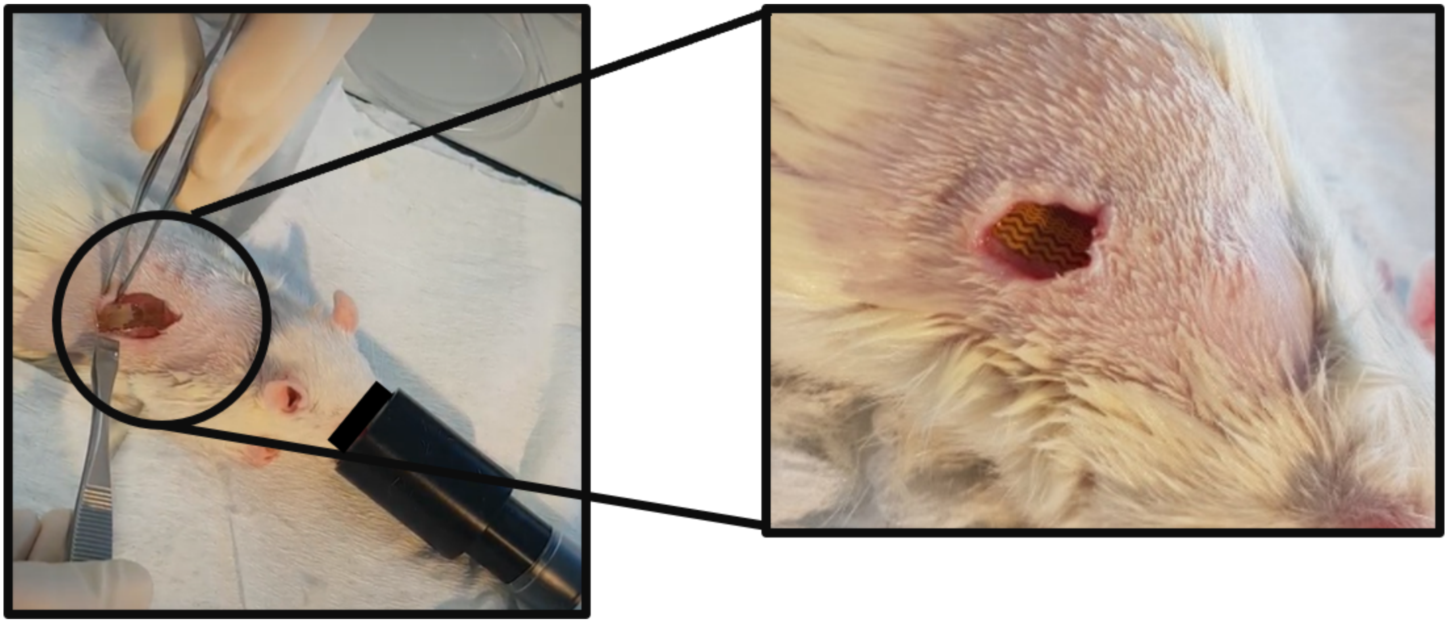

Figure S6: Surgical implantation of the TPU samples underneath the skin at the back of the rat.

Table S5: Implantation data for each sample: pure Platilon U 4201 AU, Platilon U 4201 AU with gold tracks, and medical-grade Bionate 80A. Nine samples were implanted in six female Wistar rats. Animals P1, P2 and P3 received two samples bilaterally, one material per side, providing a within-animal comparison at each of these time points. Samples with gold tracks carried laser-ablated electrode openings. "Not recorded" indicates that no qualitative observation was noted at explantation, not that the sample was unavailable for analysis.

| Animal | Sample | Side | Material | Survival (months) | Implantation | Explantation |
|---|---|---|---|---|---|---|
| P1 | 1 | left | Bionate | 2 | Uneventful | Uneventful |
| | 2 | right | Platilon | | Slightly more difficult | Uneventful |
| P2 | 5 | left | Bionate | 4 | Uneventful | Uneventful |
| | 6 | right | Platilon | | Uneventful | Uneventful |
| P4 | 4 | right | Platilon + Au | 5.5 | Large wound; uneventful | Not recorded |
| P5 | 8 | right | Platilon + Au | 6 | Uneventful | Not recorded |
| P3 | 9 | left | Bionate | 9.5 | Slightly larger wound; uneventful | Uneventful |
| | 10 | right | Platilon | | Uneventful | Not recorded |
| P6 | 12 | right | Platilon + Au | 9.5 | Uneventful | Not recorded |

Table S6: Thickness of the encapsulating connective tissue, measured for Bionate 80A (2, 4 and 9.5 months survival) and for the combined group of pure Platilon U 4201 AU and Platilon U 4201 AU with gold tracks (2, 4–6 and 9.5 months survival). Data are presented as median [first to third quartiles] of the measurement points. [a]The 4–6 month and 9.5 month Platilon groups combine pure Platilon with Platilon carrying gold tracks; the 2 month group contains pure Platilon only.

| | Bionate 80A | | | Platilon U 4201 AU[a] | | |
|---|---|---|---|---|---|---|
| Samples | 1 | 5 | 9 | 2 | 6, 4, 8 | 10, 12 |
| Implantation (months) | 2 | 4 | 9.5 | 2 | 4–6 | 9.5 |
| Number of implants | 1 | 1 | 1 | 1 | 3 | 2 |
| Measurement points ($n$) | 74 | 136 | 103 | 85 | 209 | 171 |
| Capsule thickness (µm) | 181 [147–252] | 155 [106–195] | 188 [126–318] | 111 [73–171] | 199 [139–259] | 166 [113–228] |

### S8 Elemental composition of TPU foils

Table S7: Elemental composition of TPU foils with embedded gold tracks, determined by EDX and normalized to 100%. E1: non-implanted, ultrasonicated. E2: non-implanted, as received. E3 and E4: explanted after 5.5 and 9.5 months. Values are the mean of six spectra per sample. [a]Copper was returned by the quantification routine at 0.01–0.04 wt%, which lies below the detection limit of the technique, conventionally taken as approximately 0.1 wt%; these values are not interpretable as concentrations.

| Element (wt%) | E1 | E2 | E3 | E4 |
|---|---|---|---|---|
| C | 79.88 | 79.84 | 78.44 | 78.38 |
| O | 19.42 | 19.53 | 20.72 | 20.69 |
| Si | 0.58 | 0.55 | 0.61 | 0.59 |
| S | — | — | 0.13 | 0.21 |
| Cu | below detection limit[a] | | | |
